# afMLevel: Deep learning U-Net Models for Levelling Atomic Force Microscopy Images and Movies

**DOI:** 10.64898/2026.10.19.760138

**Authors:** Maya Tekchandani, Daniel E. Rollins, Megan Westerell, Stephen D. Evans, Peter G. Adams, George R. Heath

**Affiliations:** School of Physics & Astronomy, Bragg Centre for Materials Research, University of Leeds, Leeds, UK; School of Biomedical Sciences, Astbury Centre for Structural Molecular Biology, University of Leeds, Leeds, UK

## Abstract

AFM experiments can now produce hundreds of frames per session and this rises to thousands in the case of video-rate high-speed AFM, but every image must still be levelled to remove tilt, bowing and line noise before it can be measured. Current automatic routines rely on user-chosen masks and parameters that work for some sample types but fail on others, so levelling often remains a manual, subjective bottleneck. Here, we present two U-Net models developed using machine learning for levelling AFM images without altering the sample topography. MLMask segments features so that they are excluded from conventional plane and line background fits, and MLBackground predicts the noise and artifact background directly, which is then subtracted. Both were trained on 2,001 experimental images from multiple instruments and sample types, using manually levelled images as ground truth. On 164 unseen images, MLMask routines brought 70–80% of images below a mean squared error of 0.1 relative to manual levelling, compared with 60–70% for the best multi-plane method, with a lower mean error (0.1–0.15 vs 0.15–0.3), and both models significantly outperformed fourteen classical routines (*p* < 0.01). Applied frame by frame, the models process 512 × 512 pixel HS-AFM movies at 4.7 frames s^−1^. We provide afMLevel as an openly-available package for the use of other researchers. By removing the need to choose levelling parameters image by image, this package together with its models and paired training data, provides a foundation for automated AFM analysis pipelines.

## Introduction

Advances in imaging speed [1, 2], automated acquisition [3, 4] and large-area tiling [5] mean that Atomic Force Microscopy (AFM) now yields far more data than can be processed by hand. High-Speed AFM (HS-AFM) alone reaches 1–50 frames per second (fps) [2, 6], and a single imaging session can produce hundreds of movies and many thousands of frames, enabling the structural dynamics of biological systems to be followed at the sub-molecular scale [7–9]. However, raw height images contain characteristic artefacts that must be removed before quantitative analysis: (i) line noise from z-scanner drift between consecutive scan lines, (ii) tilt from misalignment between the sample plane and the xy-scanner, and (iii) bowing from xy-scanner non-linearity. Levelling is therefore an essential step, and a bottleneck, in automated AFM analysis pipelines. Levelling typically fits polynomials to background pixels, either line by line (line fit) or across the whole image (plane fit), and subtracts them from the data. To avoid skewing these fits, features must be masked out, either by manual annotation or by height thresholding, and thresholding fails when noise is strong or feature and background heights overlap. Existing software provides iterative routines that alternate polynomial fitting with thresholding [10–13], multi-plane methods that fit regions separately [12, 14], and frequency-based removal of line noise [15]. Each suits particular image types, but none is reliable across the range of samples and artefacts encountered in practice, so a method must still be chosen and tuned for each dataset.

Deep learning offers a way to remove this choice. Machine learning is increasingly used to improve AFM acquisition and data analysis [16], and U-Net architectures [17] have been applied to the segmentation of biomolecular components [18], DNA origami [19], and proteins in HS-AFM movies [20], to general particle detection for autonomous AFM [21], and to the correction of probe-related distortions using simulated data [22].

Here we present two U-Net models designed to level AFM images without altering the topography of the sample. MLMask produces a binary feature mask that is used within conventional masked plane and line fits, and MLBackground predicts the noise background directly, which is then subtracted. We trained both models on 2,001 experimental images spanning a range of sample types and instruments, using manually levelled images as ground truth, and benchmarked them against fourteen classical routines on 164 unseen images. Both outperformed the classical methods, and we show where each succeeds and where it still fails. We also benchmark runtime on HS-AFM movies, and release the models, the afMLevel Python package [23] and the paired training data to support batch levelling and further development.

## Results

### Training Data and Ground Truth Generation

We assembled 2001 experimental AFM images from multiple instruments, manufacturers and facilities (Supplementary Figure S1), covering five sample types (membrane, protein, DNA origami, lattice and fibre) and five surface features (particles, multiple planes, strong line noise, holes and fibres; Figure 1A,B). Each image was assigned one sample type and any number of feature labels. The set deliberately over-represents challenging topographies, such as multiple planes and strong line noise (Figure 1B). Geometric augmentation and, for MLMask only, synthetic line-noise artefacts (Figure 1C) expanded the training data to approximately 10,000 images for MLMask and 6,000 for MLBackground (see Methods).

**Figure 1.**
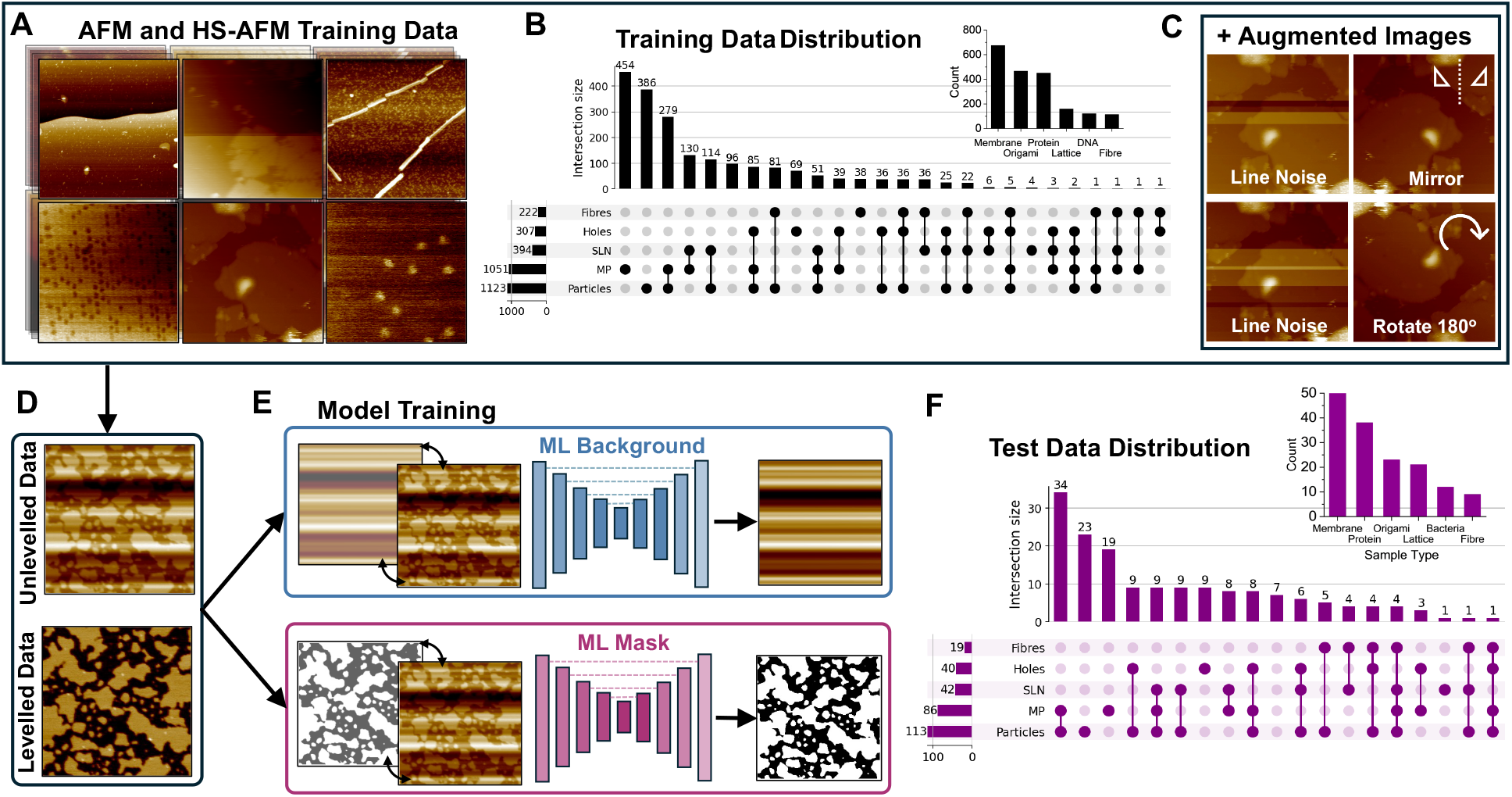
Overview of datasets, feature distributions, and model training pipelines. **(A)** Representative atomic force microscopy (AFM) and high-speed AFM (HS-AFM) images illustrating the diversity of training samples across various topographies. **(B)** Distribution of training data (*n* = 2001), showing individual sample type counts (inset bar chart) and an UpSet plot detailing the co-occurrence of topological features and artefacts. **(C)** Data augmentation strategies applied to expand training diversity, including artificial line noise injection, vertical mirroring (reflection along the y-axis), and 180^*◦*^ rotation. **(D)** Input data pairing unlevelled raw experimental scans with corresponding manually levelled ground truth references. **(E)** Schematic of the machine learning architectures consisting of the ML Background network (top) predicting continuous background variations and the ML Mask network (bottom) segmenting features of interest. **(F)** Distribution of test data (*n* = 164), displaying individual sample types (inset) and feature combinations via UpSet plot analysis.

Both models were trained using a 7-layer U-Net (Supplementary Figure S2) with full supervision using ground truth derived from manually levelled images. Each raw image was levelled manually in NanoLocz [12] using thresholding, plane fitting and line fitting. The ground-truth background was obtained by subtracting the levelled image from the raw image. The ground-truth mask was obtained by Gaussian filtering the levelled image (*σ* = 1) and applying Otsu thresholding, where the smoothing ensured that low-contrast features affected by noise were included in the mask (Figure 1D,E). We evaluated the effects of filter size, model depth, line-noise examples, batch size, activation function, preprocessing and batch normalisation to identify better performing training parameters (Supplementary Figure S3 and S4).

Performance was evaluated on a separate test set of 164 randomly selected images that were not used for training or validation (Figure 1F). Its sample-type and feature distributions resemble those of the training set, reflecting the data available, but it also includes bacterial samples, which are absent from training and therefore test the models on an unseen morphology.

Beyond training these models, the curated dataset of 2,001 raw experimental scans paired with manually generated ground-truth masks and backgrounds represents a standalone resource for the SPM community. Hosted on Hugging Face in tiff format (https://huggingface.co/datasets/Heath-AFM-Lab/afMLevel-data), the repository supports multiple programmatic access methods—including direct Python loading, on-the-fly streaming, and browser-based inspection—facilitating benchmarking and future method development in atomic force microscopy.

### Levelling Workflows

Both models share a three-stage workflow consisting of pre-processing, U-Net inference, and post-processing (Figure 2). Input images first undergo a first-order plane fit in x and y, are normalised to [0,1], and are resized to 256×256 pixels for the fixed-input networks. While MLMask achieves this via nearest-neighbour interpolation, MLBackground splits each image into multiple 256×256 arrays of alternating pixels (Supplementary Figure S5); this ensures every pixel, and consequently the fine line-by-line variation defining the noise background, is fully retained for the background calculation.

**Figure 2.**
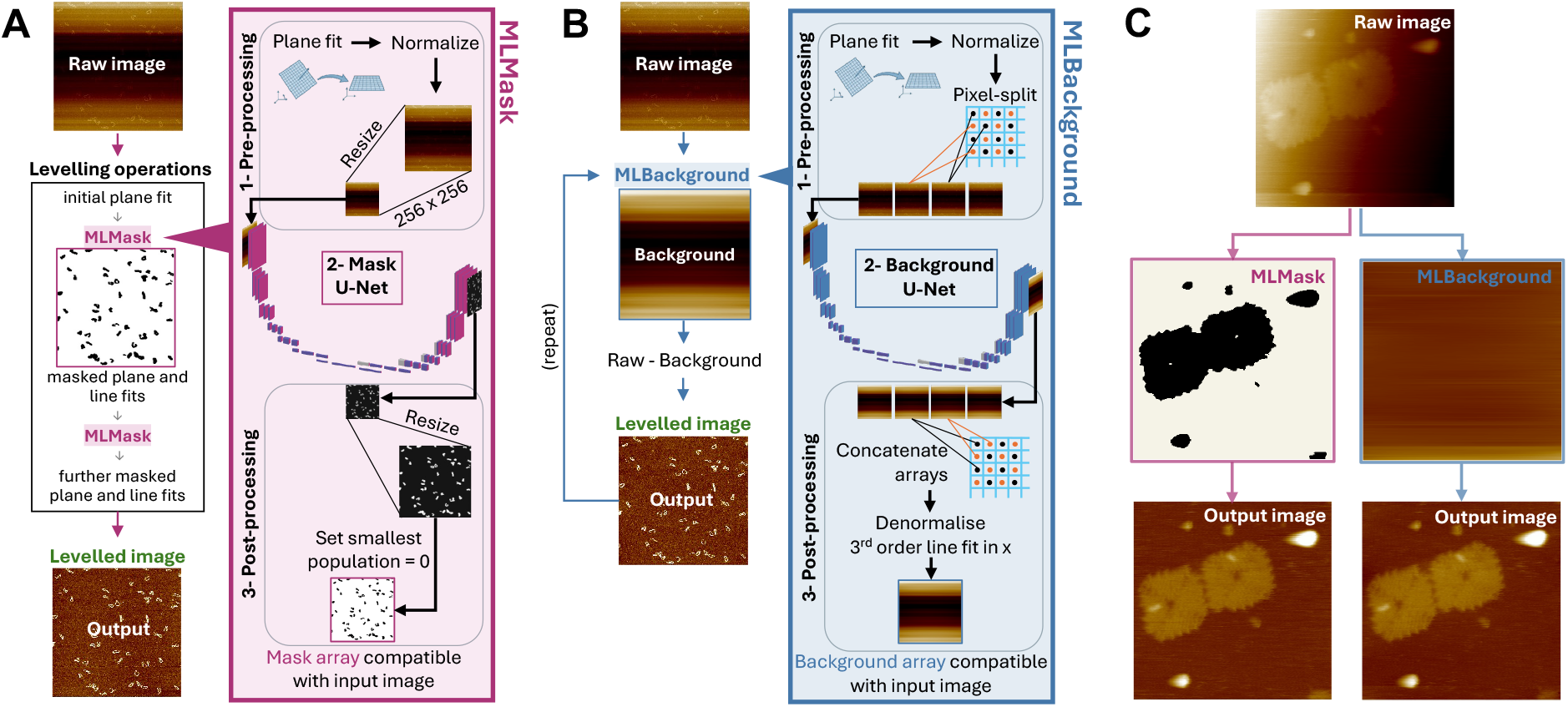
Schematic workflows of the MLMask and MLBackground machine learning pipelines. **(A)** Pipeline for the MLMask model, detailing pre-processing, U-Net feature segmentation, and iterative background and plane fitting operations. **(B)** Pipeline for the MLBackground model, incorporating pixel-split pre-processing, U-Net background prediction, concatenation, and a third-order line-wise fit in x to construct the final background estimate. **(C)** Side-by-side qualitative comparison of final output images generated by MLMask and MLBackground from a shared raw input HS-AFM frame of DNA origami.

Following pre-processing, the networks output either a binary feature mask (MLMask) or a continuous background estimate (MLBackground), after which post-processing restores the original image dimensions. For MLMask, the predicted mask is upsampled, cleaned to remove small spurious regions, and subsequently used to exclude feature pixels from plane and line fits within a conventional levelling routine. For MLBackground, the split outputs are recombined and a third-order polynomial is fitted line-by-line to the predicted background. Subtracting this fitted background rather than the raw network prediction prevents localized artefacts from distorting the underlying sample topography.

### Model Performance

We compared the ML routines with fourteen classical auto-levelling routines from NanoLocz [12] (SI Table S1), grouped as no-mask, threshold-mask and multi-plane. The no-mask method applies a plane fit with median line subtraction and excludes no pixels. Threshold-mask methods exclude features using standard deviation, fixed height, Gaussian variance or Otsu thresholds, in upper, lower or combined variants. Multiplane methods segment the image by Otsu thresholding or Sobel edge detection and apply area-weighted plane and line fits to each region. For MLMask we tested four ways of using the mask: a single-step plane-line fit (*plane-line MLMask*), an iterative routine (*iterative MLMask*), and multi-plane routines with and without an initial line fit (*multi-plane MLMask line* and *multi-plane MLMask*). For MLBackground we applied the model once, twice or three times (*MLBackground 1*×, *2*×, *3*×), using the previous output as the next input.

Figure 3A–E compares the best routine from each category on five test images that present distinct levelling challenges, combining strong line noise, multiple planes, high dynamic range, holes, and bowed backgrounds. The no-mask and threshold-mask methods often left residual artefacts because foreground pixels inadvertently entered the plane and line fits, and thresholding failed when noise was strong or feature-to-background height contrasts were low. Multi-plane methods successfully handled distinct planar regions (Figure 3D), but remained sensitive to edge-detection failures and noisy boundaries. *Iterative MLMask* segmented features robustly under high noise and yielded the best results for Figure 3A and B, where conventional masks failed to separate membrane regions and bright features from the background. However, it was limited in scenarios where masking alone is insufficient, such as strongly bowed multi-plane surfaces (Figure 3D and E). *MLBackground 2*× performed best on images dominated by background artefacts (Figure 3A, C, D, and E) because it estimates the background directly and avoids distortions from polynomial fitting; nevertheless, it struggled with the large dynamic range of Figure 3B, which excluding extreme outlier pixels partly resolved (Supplementary Figure S6).

**Figure 3.**
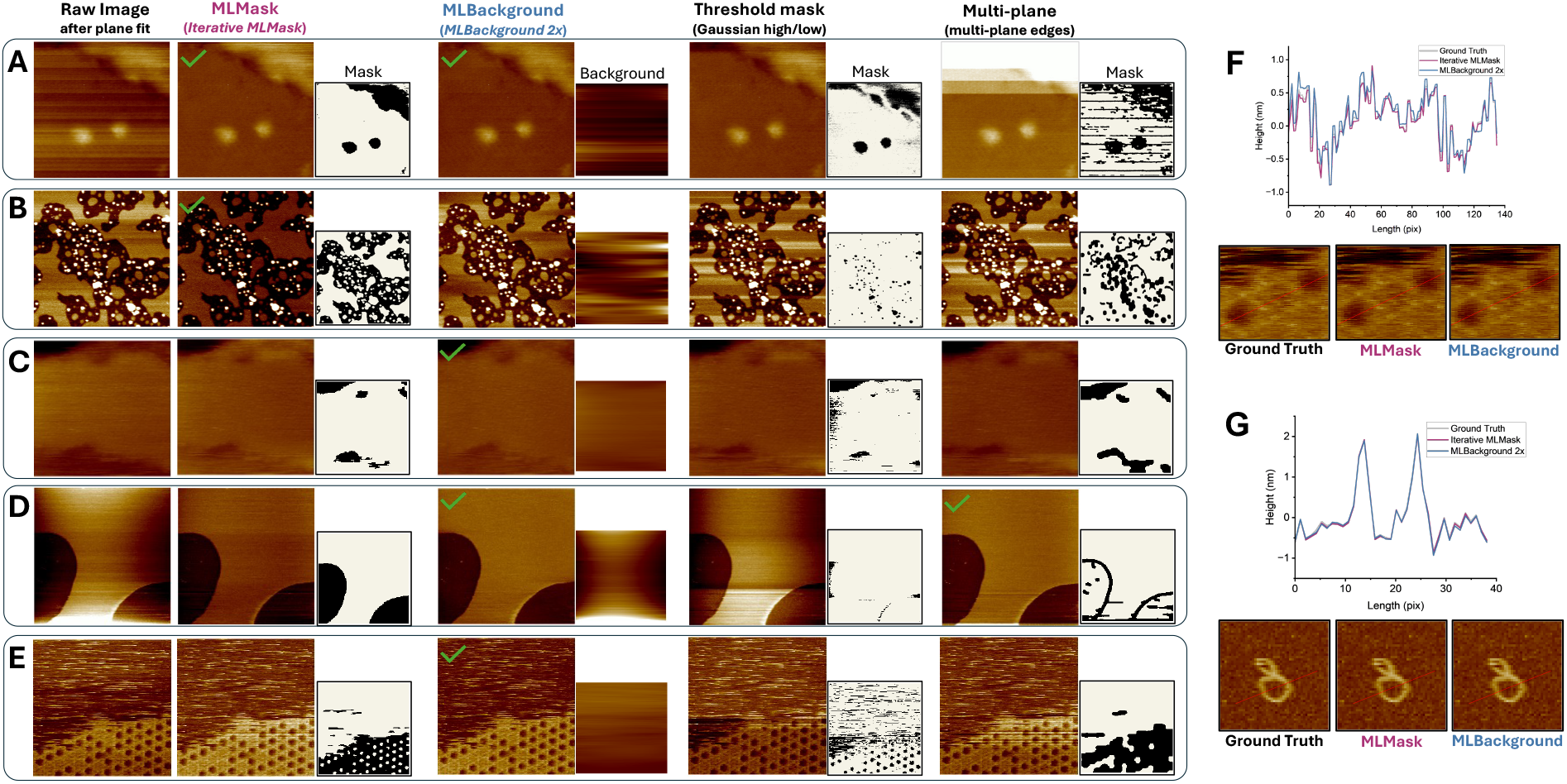
Comparison of AFM leveling performance across challenging test images and complex topographies. **(A–E)** Qualitative evaluation of five challenging AFM images. For each example, the raw input image (left) is compared against outputs from representative classical and machine learning routines: MLMask (Iterative MLMask), MLBackground (MLBackground 2x), Threshold mask (Gaussian high/low), and Multi-plane (multi-plane edges), alongside corresponding generated masks and background estimations. Green checkmarks indicate top-performing methods identified via blinded visual assessment, with multiple checkmarks denoting equivalent quality. **(F, G)** Quantitative validation using experimental ground truth scans of Annexin V **(F)** and DNA **(G)** processed with Iterative MLMask and MLBackground 2x. Corresponding cross-sectional height profiles demonstrate high quantitative fidelity and structural height preservation across complex topographies.

Cross-sectional height profiles for Annexin V and DNA (Figure 3F,G) confirm that both routines successfully preserved the local height information of the manually levelled images. To mitigate concerns that deep learning models might alter raw data through hallucination, our architectures avoid direct pixel manipulation; instead, they constrain the network to binary masks or smooth background fits before applying conventional polynomial correction. Consequently, absolute height scales and fine structural topographies remain uncompromised.

To quantify performance, we calculated the mean squared error (MSE) between each levelled output and its manually levelled ground truth following z-score normalisation, while also independently assessing each result via blinded visual inspection (categorized as good, okay, or bad). We report the mean and median MSE, the percentage of test images achieving an MSE below 0.1, and the overall success rate from visual inspection (Figure 4A,B). An MSE threshold of <0.1 was established as a practical criterion for successful leveling, isolating outputs that closely approximate the manual ground truth while minimizing susceptibility to severe outlier failures that skew mean or median metrics. Because absolute MSE remains inherently dependent on ground truth variations, visual inspection serves as a complementary, independent qualitative evaluation.

**Figure 4.**
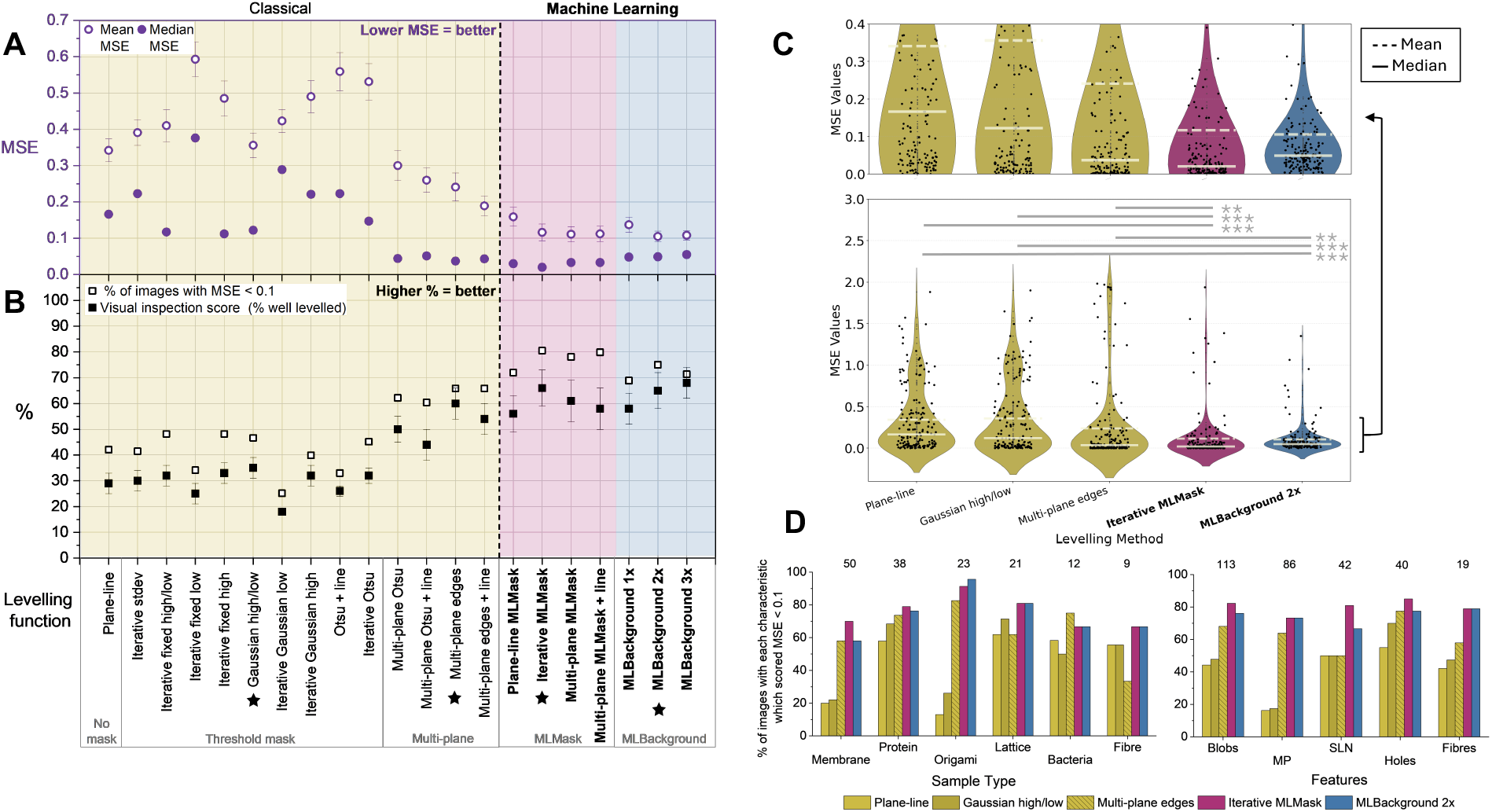
Quantitative performance evaluation of classical and machine learning levelling routines across the 164-image test dataset. **(A)** Mean and median mean squared error (MSE) for each function. **(B)** Percentage of test images achieving an MSE below 0.1 (open squares) alongside independent visual inspection scores representing the percentage of successfully levelled outputs (filled squares). Black stars denote the top-performing function within each methodological category. Error bars depict the standard error of the mean (SEM). Corresponding numerical metrics are detailed in SI Table S2. **(C)** Distribution of mean squared error (MSE) scores across the test dataset for each category of levelling method, shown as violin plots. The lower panel displays the full range of MSE values, while the upper panel provides a zoomed-in view highlighting differences at lower errors. Statistical significance between the machine learning methods and each classical baseline was assessed using Student’s *t*-test and is indicated by ** (p<0.01) or *** (p<0.001). **(D)** Success rate of each levelling method categorized by specific image characteristics. Numbers above each bar indicate the subset size (n) within the test dataset. Bar heights represent the percentage of images achieving an MSE<0.1.

Among classical routines, threshold-mask methods exhibited high variance: only 20–35% of outputs achieved successful leveling, with several routines underperforming the baseline no-mask method (30%) due to artifact introduction from sub-optimal masking. Methods targeting low-feature exclusion (*iterative fixed low, iterative Gaussian low*) performed least effectively (20%), whereas *Gaussian high/low, iterative fixed high*, and *iterative fixed high/low* achieved peak classical performance (30–35%), though occasional severe failures elevated their mean MSE. Multi-plane methods demonstrated marked improvement (median MSE <0.1; 40–60% rated good), benefiting directly from the high prevalence of multi-planar topographies (86 of 164 test images) characteristic of complex biological samples.

The MLMask routines outperformed the multi-plane methods, with 70–80% of images below MSE 0.1 and a mean MSE of 0.1–0.15, compared with 60–70% and 0.15–0.3. *Iterative MLMask* had the lowest median MSE and the highest percentage scores of any routine, although *multi-plane edges* scored higher in visual inspection than *plane-line MLMask* and *multi-plane MLMask*. The MLBackground routines performed comparably. *MLBackground 3*× had the highest visual score of all routines (~68%) but a slightly higher mean and median MSE than *MLBackground 2*×. Overall, the ML routines substantially improved upon classical methods, achieving success rates of 60–80%; the remaining failures were primarily driven by extreme topographic complexity, such as large height disparities (e.g., Figure 3B) or severe, non-uniform background artefacts that exceeded the models’ training distributions.

The MSE distributions (Figure 4C) show *iterative MLMask* clustered close to zero with the lowest median, and *MLBackground 2*× slightly broader but more symmetric with the lowest mean. The classical distributions are skewed by more frequent large failures. Both ML routines were significantly better than every classical routine, (Student’s *t*-test, *p* < 0.001 against the no-mask and threshold-mask methods, and *p* < 0.01 against the multi-plane methods). multi-plane levelling was significantly better than the no-mask and threshold-mask methods (*p* < 0.05), and the two ML routines did not differ significantly.

Across sample types, *iterative MLMask* exceeded all classical methods for membrane, protein, origami, lattice and fibre samples, whereas *multi-plane edges* was best on bacteria, which often have several height levels, and *MLBackground 2*× followed similar trends (Figure 4D). *Iterative MLMask* led on all five feature types and *MLBackground 2*× on four, equaling *multi-plane edges* for holes. Multi-plane levelling was much stronger than the other classical methods on images with multiple planes, but both ML routines improved on it slightly. In total, the ML routines outperformed the classical methods in 10 of the 11 image characteristics, and in file types they improved results on .spm and .jpk data and matched the best classical methods on .aris and .asd data (Supplementary Figure S1). Subgroup sizes are small (*n* = 9–113), so these comparisons are descriptive.

### Python Package Performance

To apply the models to raw images and HS-AFM videos, we released the Python package afMLevel [23], which handles pre-processing, inference and output generation, and tested it as a plugin for the playNano HS-AFM analysis platform [24]. *Iterative MLMask* and *MLBackground 2*× were applied frame by frame to HS-AFM videos (Figure 5A) on a laptop with a GPU (see Methods) and timed against the classical *iterative fixed high* routine.

**Figure 5.**
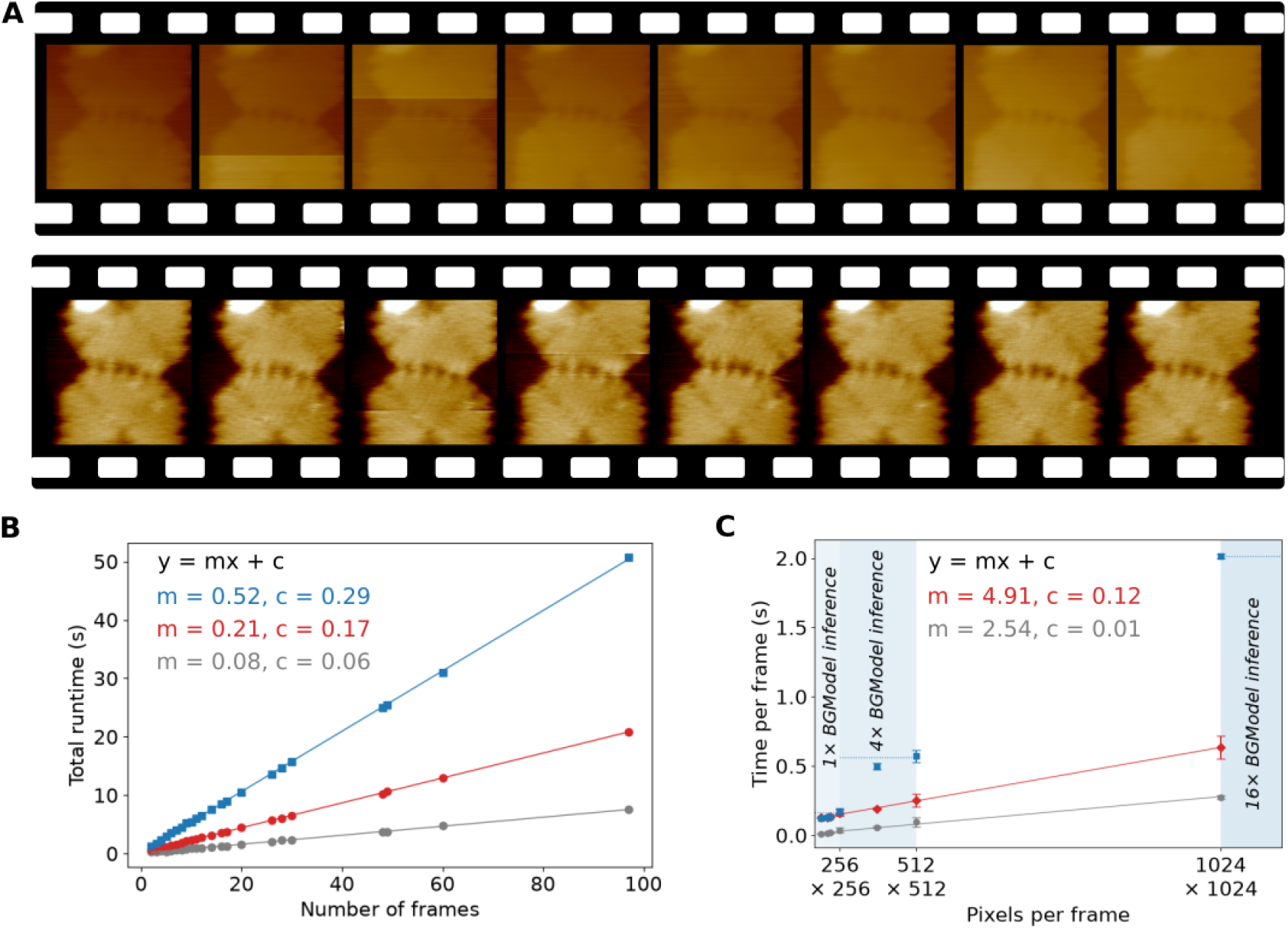
Analysis of the Python package performance on a range of HS-AFM videos. (A) Example frames from a HS-AFM video of a DNA origami structure showing the raw unprocessed data (above) and the data processed with *MLBackground 2* × model through playNano (below). (B) Plot of the runtime for the *Iterative MLMask* routine (red), *MLBackground 2* × model (blue) compared with a classical *iterative fixed high* routine (grey) against the number of frames in HS-AFM video. A linear fit (y = mx +c) was applied to the datapoints from each routine and the fit parameters are shown on the graph. (C) Plot of time taken to process a frame of a HS-AFM video (total time/ number-of-frames) against the resolution of the videos, in pixels per frame, showing the linear relation between these for the *Iterative MLMask* routine and the classical *iterative fixed high* method and the stepped increase for the *MLbackground 2* × routine. Again, the fit parameters are shown.

Total processing time scaled linearly with the number of frames for all routines (Figure 5B). For 512× 512 pixel images, *iterative MLMask* ran at 4.7 frames s^−1^, compared with 13.0 frames s^−1^ for the classical routine, and both scaled approximately linearly with the number of pixels (Figure 5C). *MLBackground* scales in steps instead, because each additional 256 × 256 tile requires another U-Net inference, so its time per frame is similar across resolutions that map to the same number of tiles (Figure 5C). For a 97-frame, 256 × 256 pixel video, the classical routine took 2.58 s, *iterative MLMask* 13.95 s and two passes of MLBackground 14.32 s. The ML routines are therefore roughly 3–5 times slower than the classical routine, but processing times remain practical for research use and substantially shorter than manual levelling.

## Discussion

afMLevel and its models demonstrate that machine-learning-based levelling provides a more generalisable, bias-free alternative to conventional AFM workflows. Unlike classical approaches requiring manual selection of masking and fitting parameters, both models adapt to diverse sample types and noise conditions. While occasionally serving as the sole successful approach for complex images, their primary advantage lies in broad robustness, representing an important step toward fully automated AFM image processing.

Despite these advantages, both models retain limitations. MLMask remains dependent on subsequent polynomial fitting and is restricted by binary height thresholding, preventing accurate masking of images featuring both high and low structures. MLBackground occasionally mistakes real horizontal features for background noise, rendering it unsuitable for fibers oriented along the scan axis (Supplementary Figure S8), and struggles with high-dynamic-range images where extreme height values distort normalisation. Although excluding values beyond 4 standard deviations on a second pass resolves this for some images, it degrades performance on standard topographies (Supplementary Figure S6), yielding a worse median MSE; thus, standard processing is retained in the general package. Users are encouraged to explore multiple routines within afMLevel to determine the optimal output for specific samples. Our architectural choices carefully balance performance and computational cost. Denoising U-Nets often reconstruct full images, presenting challenges in preserving 3D topographic structural and height information despite architectural improvements [25, 26]. Following the approach of Zhang et al. [27] with DnCNN, MLBackground avoids full image reconstruction by predicting only the noise background; applying a third-order line fit then removes local pixel artefacts without altering sample topography. Similarly, while segmentation U-Nets can suffer from resolution loss, motivating nested architectures like U-Net++ [28, 29], these increase runtime (Figure 5).

Because levelling relies primarily on dominant foreground structures, a standard U-Net provides a sufficient balance of accuracy and efficiency for both models.

Further improvements will stem from architectural refinements and expanded training data. To address unconventional noise patterns under-represented in training data, adapting image priors (such as bilateral filter residuals or gradient information, as explored for optical images by Soniya et al. [30]) could enhance MLBackground. Additionally, physically motivated synthetic AFM datasets and physical noise models can reproduce realistic artefacts for augmentation and distortion removal, as demonstrated by Ruben et al. [21] and Kocur et al. [22]. Unlike those end-to-end reconstruction approaches, our separation of noise estimation from levelling explicitly preserves feature height during post-processing.

A key bottleneck remains the scarcity of diverse training datasets siloed across individual laboratories. To support the community, we release our paired training data, sourced from our facilities and external online repositories [31–40], alongside the workflows. Future versions of afMLevel could also integrate uncertainty estimation to flag unreliable automated outputs for manual inspection.

In conclusion, afMLevel offers significant improvements over classical auto-levelling functions, dramatically accelerating processing times particularly for video AFM data. MLMask is recommended for strong line noise, while MLBackground is suited for artifact-free background visualization. All workflows are publicly available on GitHub [23] and the trained models on Hugging Face [41, 42].

## Methods

### Data Sources and Training Dataset Preparation

Raw AFM data were sourced from experiments conducted at the University of Leeds (n = 1452), comprising datasets acquired between 2010 and 2025. Additional raw AFM data were obtained from Simon Scheuring (Weill Cornell Medicine), Abimbola Feyisara Olulana (University of Sheffield) and from publicly available online repositories [31, 33, 35, 37–40]. To reduce over representation of individual datasets, images extracted from HS-AFM movies were limited to a maximum of 20 frames per movie. Ground-truth levelled images were generated through manual levelling using plane fitting, line fitting and multi-plane fitting with user selected manual thresholds and when beneficial automatic edge detection. The resulting paired dataset, consisting of raw input images and corresponding ground-truth levelled images, has been made openly available (https://huggingface.co/datasets/Heath-AFM-Lab/afMLevel-data)[43] to support reproducibility and future machine-learning development within the AFM community.

When labelling the images to characterise them based on the features present, the following definitions were applied: ’Particles’ refer to any sort of individual particles with a raised height relative to the surface they are on; ’Multiple planes’ refer to at least two flat regions of different heights; ’Strong line noise’ refers to images with dominant stripe artefacts, equal to or greater than the height range of the features; ’Holes’ refer to small regions of lower height relative to the surface, and ’Fibres’ refer to long fibrous features.

The images in each subset were augmented to multiply the size of the dataset by 3 for MLBackground and by 5 for MLMask (including 2 images with artificial line noise per image). This gave 8005:2000 training:validation images for the mask model and 4803:1200 training:validation images for the background model. Augmentation consisted of two geometric transformations (180 degree rotation and horizontal reflection), chosen to preserve the direction of the horizontal line noise. For MLMask only, because it improved the loss convergence, additional synthetic line-noise artefacts were introduced by adding randomly generated stripe patterns with constrained number, thickness, and amplitude to mimic experimentally observed noise. The artificial line noise was created by adding an array of randomly generated stripes to the image, with constraints on the amount (between 1 and 3), the stripe thickness (between 5 and 50 pixels), and the height (plus or minus a value between the image minimum and ¼ image maximum), so that it resembled realistic line noise.

### Model Training

The mask and background U-Net models were created and trained in Python using functions from the PyTorch library. The model architecture is shown in Supplementary Figure S2, adapted from the original U-Net architecture [17]. Training parameters were varied methodically and the loss curves compared to choose values which minimised the loss for the validation data. These loss curves are shown in Supplementary Figure S3 (mask model) and Supplementary Figure S4 (background model). The loss functions used were the sum of the BCE loss and Dice loss for the mask model (equations 1. 2), and MSE for the background model (equation 3).

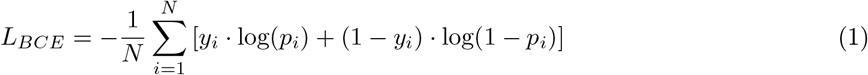

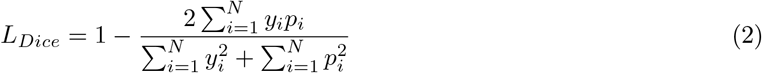

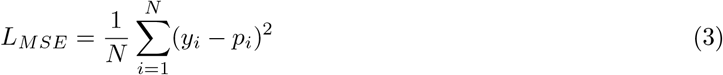

BCE provides a pixel-wise comparison by calculating the difference between the predicted value for each pixel and the groundtruth value of either 0 or 1. Dice loss measures the global similarity, with the numerator representing the intersection of the two images and the denominator normalising for their sizes. This combined loss function has been shown to perform well in segmentation tasks compared to other loss functions [44]. Like BCE, MSE is a pixel-wise comparison, but it is more suited to images with a continuous range of values, such as the background array, whereas BCE was made specifically for binary classification tasks.

The final optimised models had 7 layers and used filters of 7×7 pixels (mask) or 9×9 pixels (background). Each downsampling path layer consisted of two convolutional blocks, each followed by Batch Normalisation (BN) and activation using the Rectified Linear Unit (ReLU) function, and then max pooling to reduce the dimensions for improved efficiency. The upsampling path used up-convolution between layers to return the feature maps to the original dimensions. In each layer, skip connections merged the corresponding feature maps from the downsampling path with the upsampling feature maps to retain spatial details lost during downsampling. The gradient descent algorithm used during training to update the weights (filter values) was the Adam Optimiser [45], due to its speed and success in image processing tasks, with a learning rate of 0.0005 to provide a smooth loss reduction. The models were trained using an initial data set of 2,001 unique raw AFM images, which was randomly split at 80:20 into training and validation subsets.

### Model Testing and Scoring

All model testing and scoring was performed in MATLAB to allow direct comparison with pre-existing auto-levelling functions in the MATLAB software, Nanolocz [12]. Four MLMask levelling routines (MLMask, iterative MLMask, multi-plane MLMask and multi-plane MLMask + line) and three MLBackground routines (MLBackground 1×, MLBackground 2× and MLBackground 3×), detailed in SI table S1, were created in MATLAB to perform levelling using the models. Within these routines, the models themselves (red and blue box, respectively, in Figure 2) were applied by running python inference scripts within MATLAB workflows using the MATLAB Python interpreter. Fourteen classical MATLAB levelling routines were tested alongside the ML levelling routines (SI table S1).

The levelling functions were applied to each image in the test dataset (Figure 1B), and the MSE between the auto-levelled image and the corresponding ground truth (manually levelled image) was used as the main metric to score and compare the levelling methods. Before calculating the MSE, all images were pre-processed by z-scale normalisation, shifting the mean pixel intensity to 0 and standard deviation to 1. This was done to prevent differences in intensity scales between images from affecting the MSE score.

### Visual Inspection

A visual inspection of levelling performance was performed by applying the levelling functions to each image in the test dataset, and viewing the anonymised output (levelled image) for each function side-by-side in a randomised order. The levelling on each image was then scored as ’good’, ’okay’, or ’bad’. The categories were defined as follows: ’good’ if the image looked perfectly levelled, ’okay’ if it was almost perfectly levelled but there were some small flaws (e.g. slight tilt or one small line remaining), and ’bad’ if there were obvious artefacts in the image. This was repeated by three authors to obtain averaged results. A demonstration of the visual inspection scoring is given in Figure S7.

### Performance Evaluation of afMLevel Workflows

The performance of the levelling workflows was evaluated using a custom Python timing script (https://gist.github.com/derollins/9c5ec2dfd15dfe209a4733707c9915a8) built around the afMLevel Python package [23], integrated as a plugin within the playNano high-speed AFM analysis platform [24]. The script applied predefined processing routines to AFM image stacks and recorded total execution times under controlled conditions. All processing was performed on a laptop with an Intel Core Ultra 7 165H processor, 32 GB RAM, and an NVIDIA RTX 500-series laptop GPU, running a 64-bit Windows 11 operating system.

Input data consisted of high-speed AFM (HS-AFM) image stacks loaded via playNano from several publicly available datasets [31, 37–40]. For each dataset, both trace and retrace height channels were processed independently if present.

Processing workflows were implemented using the playNano processing pipeline interface to apply sequential processing steps. The evaluated routines included MLBackground applied one, two, and three times; MLMask applied in both the ’MLMask’ and ’iterative MLMask’ routines; and a classical comparison using the ’iterative 1 nm high’ routine from the Python Nanolocz implementation [46]. Each routine was repeated three times per dataset, and timing results were recorded for each repeat. For each processing routine, a linear model was fitted to the mean processing time as a function of the number of frames using ordinary least-squares regression. The resulting slope was used as the estimate of the per-frame computational cost.

No fine-grained profiling of individual pipeline steps was performed; timing measurements therefore reflect total end-to-end execution time for each processing workflow.

## Acknowledgements

The authors thank Varun Gupta, Bashra Mahamed and Yueqi Wang for providing AFM data and feedback on the implementation of afMLevel models. We thank Chalmers Chau, Simon Connell, Simon Scheuring and Abimbola Feyisara Olulana for providing additional training data. Maya Tekchandani acknowledges studentships funded by the Engineering and Physical Sciences Research Council and the Biotechnology and Biological Sciences Research Council. Daniel E. Rollins and George R. Heath acknowledge funding from the Engineering and Physical Science Research Council grant EP/W034735/1.

## Code and Model Availability

The afMLevel software package is available on GitHub at https://github.com/mayatek1/afMlevel and is distributed through PyPI (https://pypi.org/project/afmlevel/ or pip install afmlevel). The repository contains all code required to apply the afMLevel models to NumPy arrays of AFM image data, together with example notebooks demonstrating how raw AFM data can be loaded with external packages and then processed with afMLevel. The script used to time the processing pipelines is available as a GitHub Gist here: https://gist.github.com/derollins/9c5ec2dfd15dfe209a4733707c9915a8

The model training and testing scripts will also be made publicly available through the afMLevel GitHub repository at the time of publication. A versioned release corresponding to this manuscript will be archived in Zenodo and assigned a DOI. The trained U-Net models used by afMLevel are automatically downloaded when required during execution or publicly available through the Heath-AFM-Lab organisation on Hugging Face (https://huggingface.co/Heath-AFM-Lab) [41, 42].

## Data Availability

No new experimental data has been generated in this study. The AFM datasets used for model training and testing are publicly available through the Heath-AFM-Lab organisation on Hugging Face (https://huggingface.co/datasets/Heath-AFM-Lab/afMLevel-data)[43]. All datasets used in this study are referenced in the manuscript and are available under a CC BY 4.0 licence.

## Conflict of Interest

The authors declare no conflict of interest.

## Notes

https://huggingface.co/datasets/Heath-AFM-Lab/afMLevel-data

https://github.com/mayatek1/afMlevel

### Competing Interest Statement

The authors have declared no competing interest.

