## Supplementary Information for "afMLevel: Deep learning U-Net Models for Levelling Atomic Force Microscopy Images and Movies"

---

This Supplementary Information contains 2 Supplementary Tables and 8 Supplementary Figures.

| Category | Function name | Mask | Operations |
|---|---|---|---|
| No mask | Plane-line | none | 1st order plane fit (x, y) → median line subtraction |
| Threshold mask | Iterative stdev | > mean + 2 std. dev<br>< mean - 2 std. dev | Iterative routine:<br>1st order plane fit (x, y) → 2×[mask → 1st order plane fit (x, y)] → mask → median line subtraction → 1st order plane fit (x) → mask → median line subtraction → 2nd order plane fit (x) |
| | Iterative fixed high/low | > mean + 1 nm<br>< mean - 1 nm | Iterative routine |
| | Iterative fixed low | < mean - 1 nm | Iterative routine |
| | Iterative fixed high | > mean + 1 nm | Iterative routine |
| | Gaussian high / low | > largest peak + 1.5 $\sigma^2$<br>< largest peak - 1.5 $\sigma^2$ | 1st order plane fit (x, y) → median line subtraction → Gaussian fit to histogram of pixel values → mask → 1st order plane fit (x, y) → median line subtraction |
| | Iterative Gaussian low | < largest peak - 1.5 $\sigma^2$ | Iterative routine Gauss:<br>2×[1st order plane fit (x, y) → median line subtraction → Gaussian fit to histogram of pixel values → mask] → 1st order plane fit (x, y) → median line subtraction |
| | Iterative Gaussian high | > largest peak + 1.5 $\sigma^2$ | Iterative routine Gauss |
| | Iterative Otsu | > Otsu threshold | Iterative routine |
| | Otsu + line | > Otsu threshold | 1st order line fit (x) → mask → 2nd order line fit (x) |
| Multi plane | Multi-plane Otsu | Otsu threshold edges | Iterative multi-plane routine:<br>1st order plane fit (x, y) → 3×[mask → weighted 2nd order plane fit (x, y)] → weighted median line subtraction → mask → weighted 2nd order plane fit (x, y) → weighted median line subtraction |
| | Multi-plane Otsu + line | Otsu threshold edges | Iterative multi-plane routine<br>+ median line subtraction after step 1 |

| Category | Function name | Mask | Operations |
|---|---|---|---|
| | Multi-plane edges | Sobel edges | Iterative multi-plane routine |
| | Multi-plane edges + line | Sobel edges | Iterative multi-plane routine + median line subtraction after step 1 |
| MLMask | Plane-line MLMask | MLMask | 1st order plane fit (x, y) → mask → median line subtraction → 1st order plane fit (x, y) |
| | Iterative MLMask | MLMask | Iterative routine |
| | Multi-plane MLMask | MLMask edges | Iterative multi-plane routine |
| | Multi-plane MLMask + line | MLMask edges | Iterative multi-plane routine + median line subtraction after step 1 |
| MLBackground | MLBackground 1× | – | MLBackground |
| | MLBackground 2× | – | Apply MLBackground 2× |
| | MLBackground 3× | – | Apply MLBackground 3× |

Table S1. The operations applied to the raw AFM image in each auto-levelling routine. The mask sets all masked pixels to 0 so they are not used in the calculation of the following plane or line fits. The multi-plane functions use the threshold operations to detect and mask out the edges of features, so that planes of different heights can have polynomials calculated separately, and then a weighted average of the polynomials for each region is applied to the image.

| | **Levelling function** | Mean MSE | Median MSE | MSE <0.1 (%) | Mean visual inspection score (%) |
|---|---|---|---|---|---|
| No mask | Plane-line* | 0.34 ± 0.03 | 0.17 | 42 | 29 ± 4 |
| Threshold mask | Iterative stdev | 0.39 ± 0.04 | 0.22 | 41 | 30 ± 4 |
| | Iterative fixed high/ low | 0.41 ± 0.04 | 0.12 | 48 | 32 ± 4 |
| | Iterative fixed low | 0.59 ± 0.05 | 0.38 | 34 | 25 ± 4 |
| | Iterative fixed high | 0.49 ± 0.05 | 0.11 | 48 | 33 ± 4 |
| | Gaussian high/ low* | 0.36 ± 0.03 | 0.12 | 47 | 35 ± 4 |
| | Iterative Gaussian low | 0.42 ±0.03 | 0.29 | 25 | 18 ± 1 |
| | Iterative Gaussian high | 0.49 ± 0.04 | 0.22 | 40 | 32 ± 4 |
| | Otsu + line | 0.56 ±0.05 | 0.22 | 33 | 26 ± 2 |
| | Iterative Otsu | 0.5 ± 0.1 | 0.15 | 45 | 32 ± 3 |
| Multi plane | Multi-plane Otsu | 0.30 ± 0.04 | 0.04 | 62 | 50 ± 5 |
| | Multi-plane Otsu + line | 0.26 ± 0.03 | 0.05 | 60 | 54 ± 6 |
| | Multi-plane edges* | 0.24 ± 0.04 | 0.04 | 66 | 60 ± 6 |
| | Multi-plane edges + line | 0.19 ± 0.03 | 0.04 | 66 | 54 ± 6 |
| MLMask | Plane-line MlMask | 0.16 ± 0.03 | 0.03 | 72 | 63 ± 4 |
| | Iterative MLMask* | 0.12 ± 0.02 | 0.02 | 80 | 66 ± 7 |
| | Multi-plane MLMask | 0.11 ± 0.02 | 0.03 | 78 | 61 ± 8 |
| | Multi-plane MLMask + line | 0.11 ± 0.02 | 0.03 | 80 | 58 ± 8 |
| MLBackground | MLBackground 1× | 0.14 ± 0.02 | 0.05 | 69 | 58 ± 6 |
| | MLBackground 2×* | 0.11 ± 0.01 | 0.05 | 75 | 65 ± 7 |
| | MLBackground 3× | 0.11 ± 0.01 | 0.06 | 71 | 68 ± 6 |

Table S2. Values corresponding to the graph in figure 4, showing the performance of the levelling functions on the 164 test images. The errors shown are the standard error in the mean, this being between the 164 MSE scores for mean MSE and between three repeats of scoring by visual inspection.

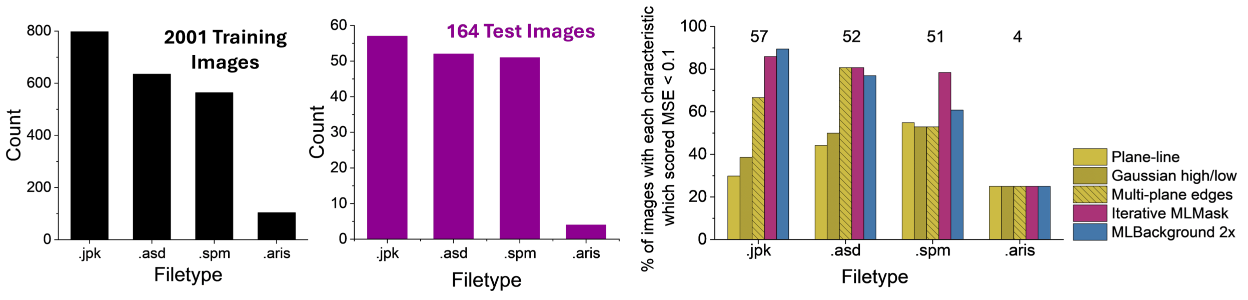


Figure S1. **A** Distribution of file types in the training (left) and test (right) datasets. The file type (.spm, .jpk, .asd, or .aris) correspond to several AFM instruments including various MultiModes (Bruker), Dimension FastScans (Bruker), JPK NanoWizards (Bruker), a JPK NanoRacer HS-AFM (Bruker), various SS-NEX HS-AFMs (RIBM) and Cypher VRS1250 (Oxford Instruments). **B** Performance of each levelling method across different image file types. Numbers above each bar indicate the number of images in the test dataset with the corresponding file type. Bar height represents the percentage of images achieving an MSE of <0.1 for each method..

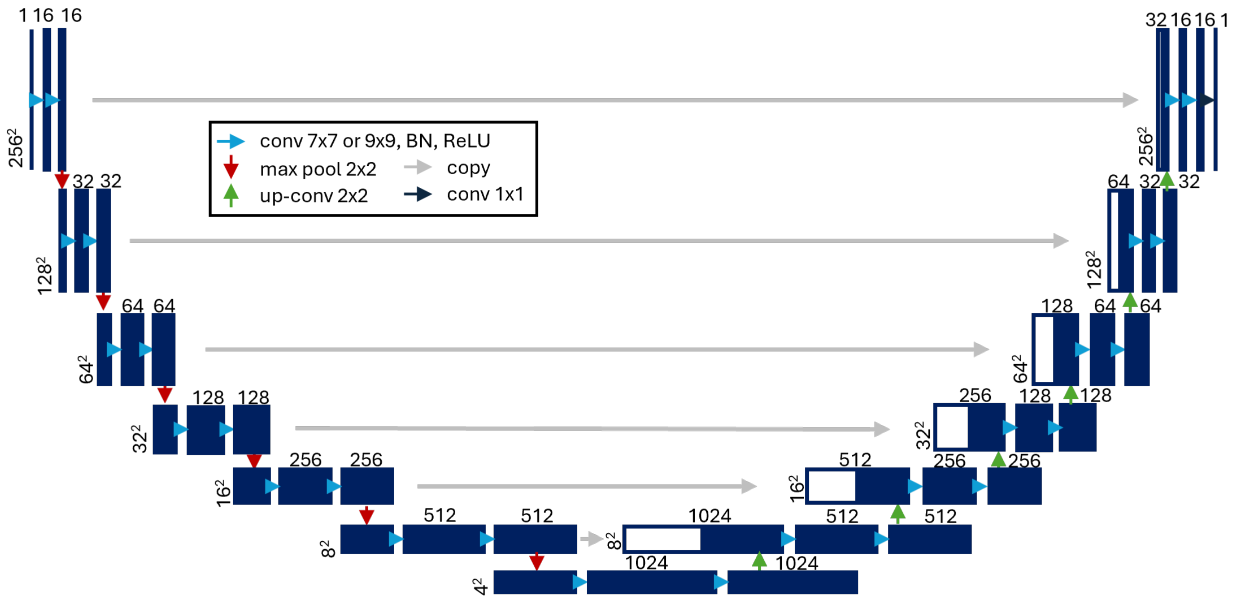


Figure S2. The 7-layer U-Net architecture used for the MLBackground and MLMask models. BN = Batch Normalisation. The first tile represents the input image array and the other tiles represent the feature maps produced by the operations indicated by the different coloured arrows (blue - convolution; red - max pooling; green - upsampling; grey - skip connections). The numbers above and to the left of the feature maps represent their number of channels and x, y dimensions respectively.

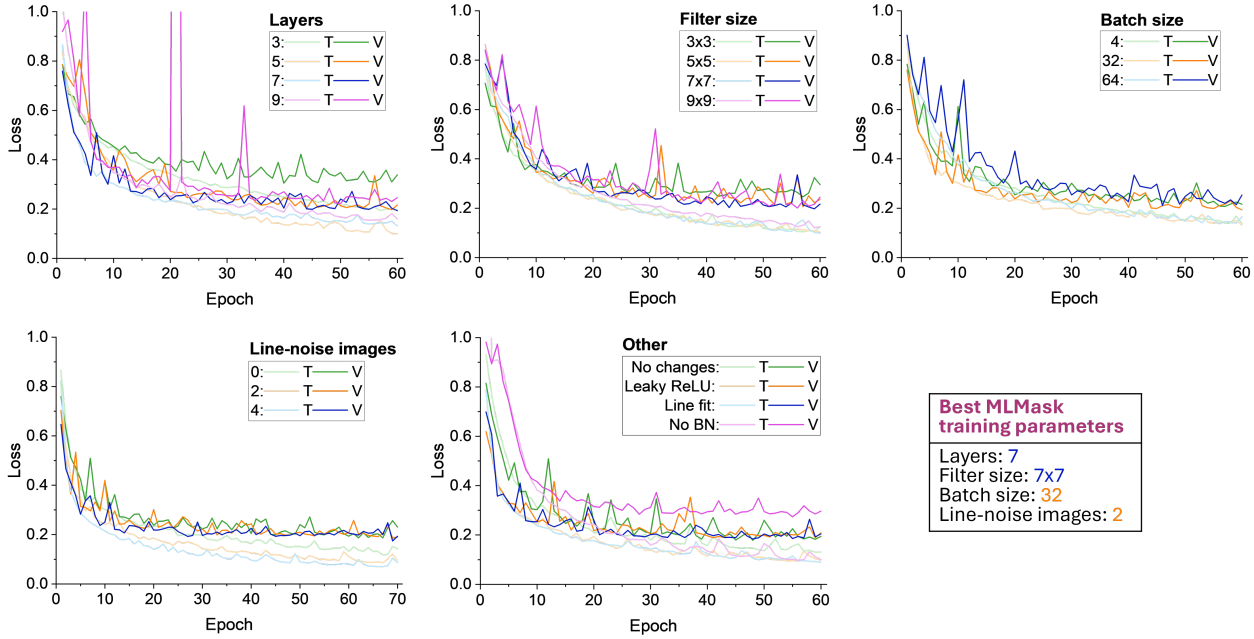


Figure S3. Comparison of MLMask loss curves to determine optimal training parameters. T = training loss curve; V = validation loss curve. For each parameter test, all other parameters were kept constant: the filter size test used 5 layers, 0 line images and batch size 32; the model complexity (layers) test used filter size $7 \times 7$, 0 line images and batch size 32; the line-noise test used filter size $7 \times 7$, 7 layers and batch size 32; the batch size test used filter size $7 \times 7$, 7 layers and 0 line images. The extra changes test used the best parameters from the preceding testing (filter size $7 \times 7$, 7 layers, 2 line images and batch size 32) with a different activation function (Leaky ReLU rather than ReLU), additional preprocessing (initial median line subtract from the raw training images), and with batch normalisation (BN) removed.

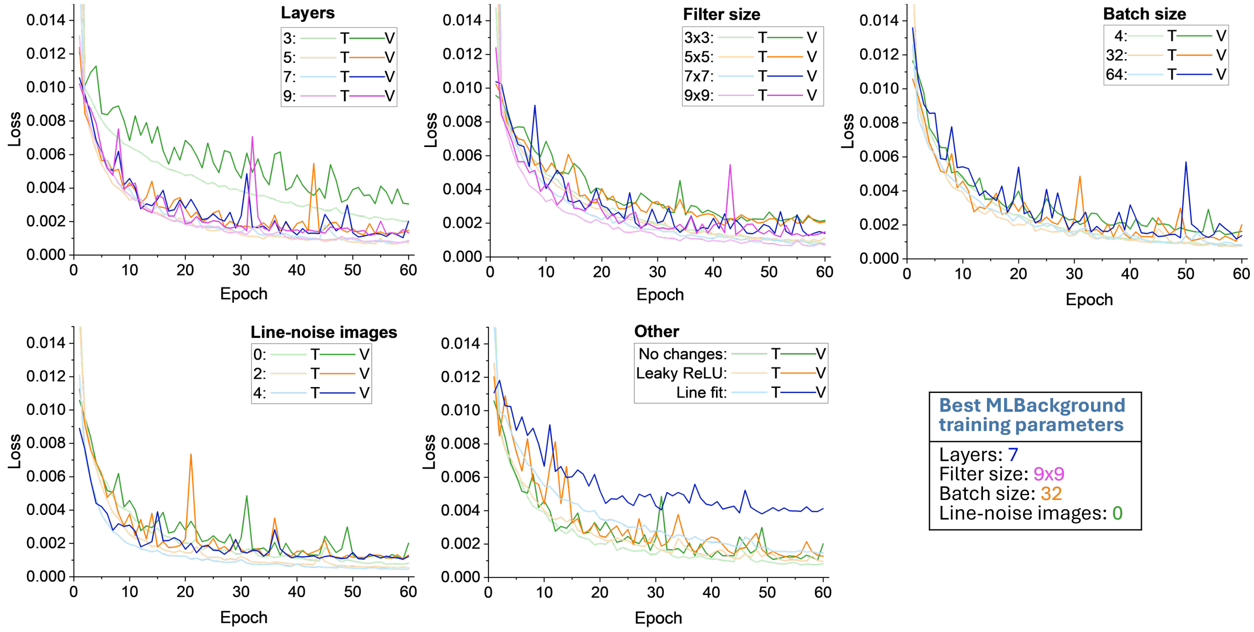


Figure S4. Comparison of MLBackground loss curves to determine optimal training parameters. T = training loss curve; V = validation loss curve. For each parameter test, all other parameters were kept constant: the filter size test used 5 layers, 0 line images and batch size 32; the model complexity (layers) test used filter size 9 × 9, 0 line images and batch size 32; the line-noise test used filter size 9 × 9, 7 layers and batch size 32; the batch size test used filter size 9 × 9, 7 layers and 0 line images. The extra changes test used the best parameters from the preceding testing (filter size 9 × 9, 7 layers, 0 line images and batch size 32) with a different activation function (Leaky ReLU rather than ReLU) and additional preprocessing (initial median line subtract from the raw training images). Running without batch normalisation gave loss values too high to be visible in the plot.

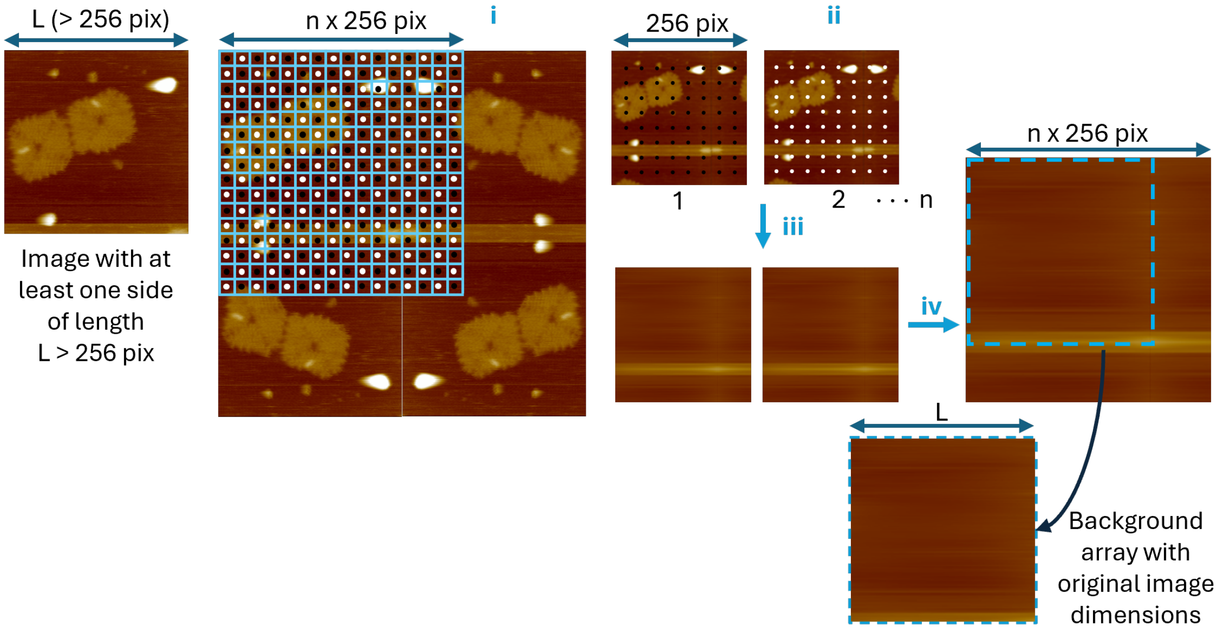


Figure S5. The steps used to resize arrays within the background model, using a pixel-split method. (i) Pad edges by mirroring array and crop to next multiple (n) of 256; (ii) Split cropped section into n x (256,256) arrays by taking every nth pixel, starting from pixel 1 to n respectively ; (iii) apply the U-Net each (256,256) array and (iv) Concatenate and crop back to original image dimensions. This shows the application for images with dimensions greater than 256 $\times$ 256. For images smaller than (256,256) pix, the array is padded by mirroring and cropped to (256,256), so no splitting or concatenation is needed (i.e. n $=$ 1).

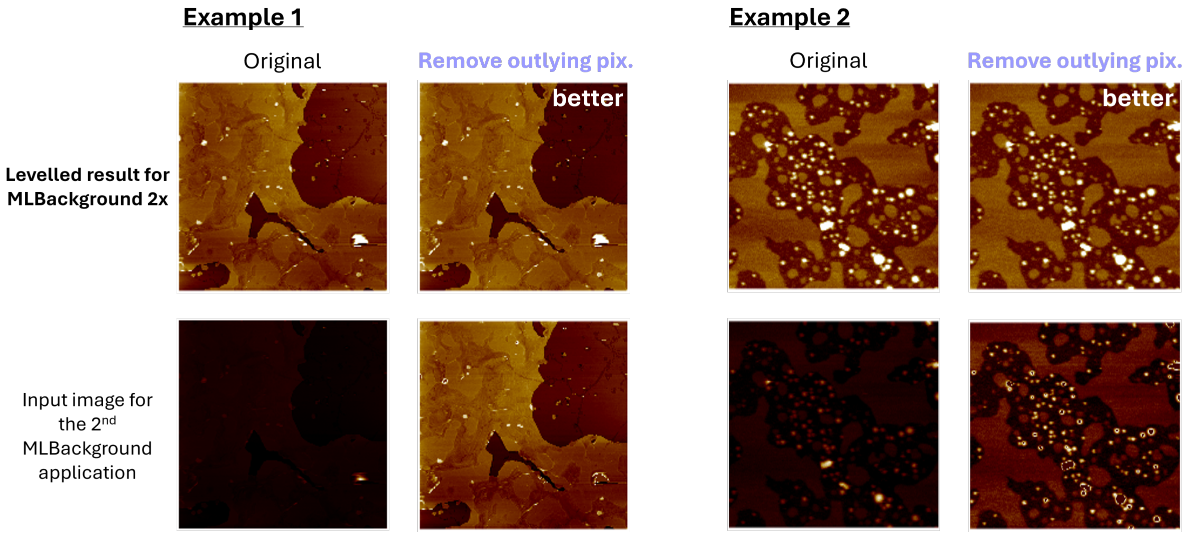


Figure S6. Demonstration of improved levelling of images with high features by setting outlying pixels (outside 4 standard deviations from the mean) to the mean value before the second application of MLBackground in *MLBackground 2*$\times$.

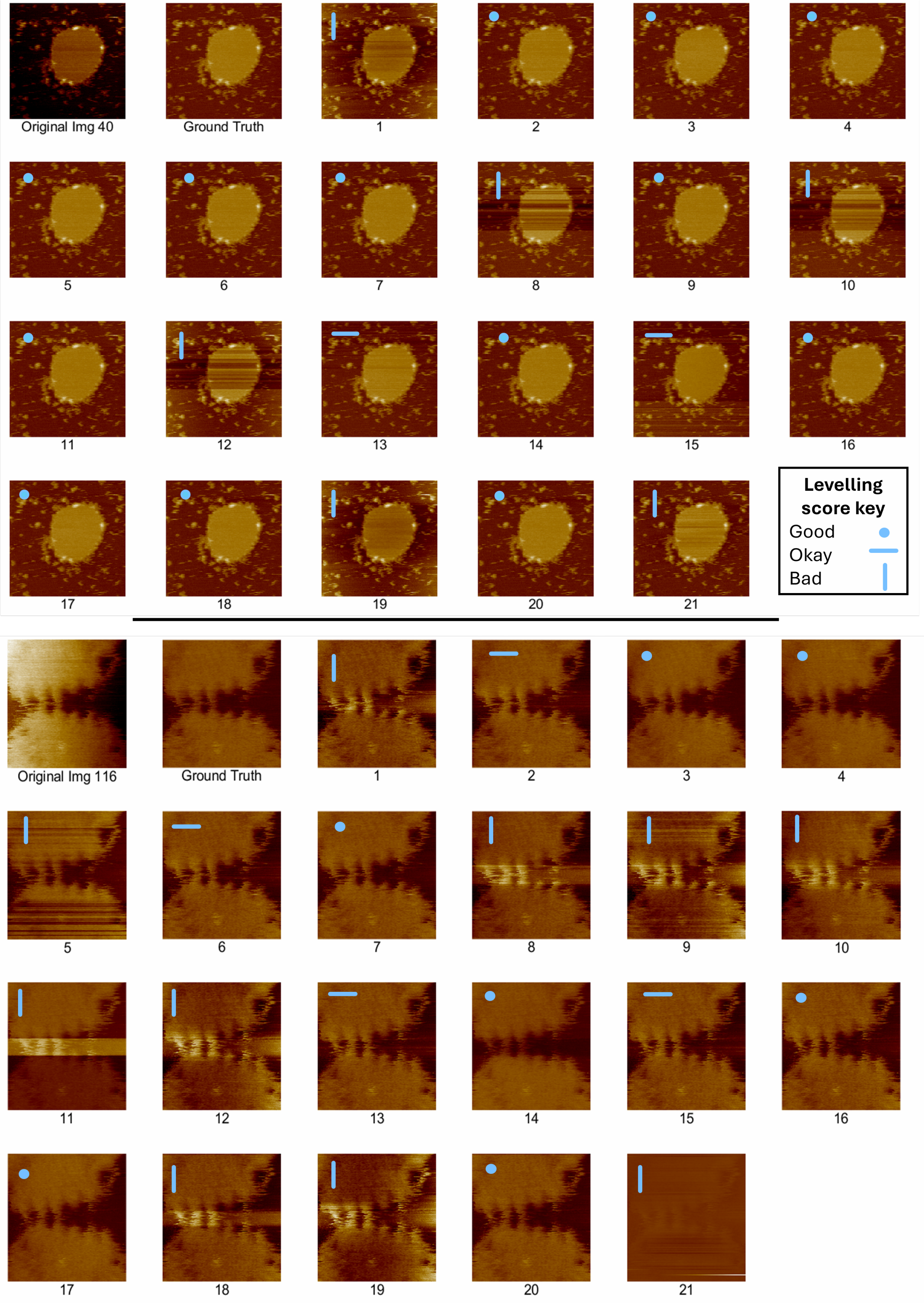


Figure S7. Example of how the visual inspection was carried out on two images from the test dataset. For each image, the raw image and ground truth are displayed alongside the levelled results from each levelling function, anonymously labelled 1-21. The results were then scored as 'good', 'okay' or 'bad'.

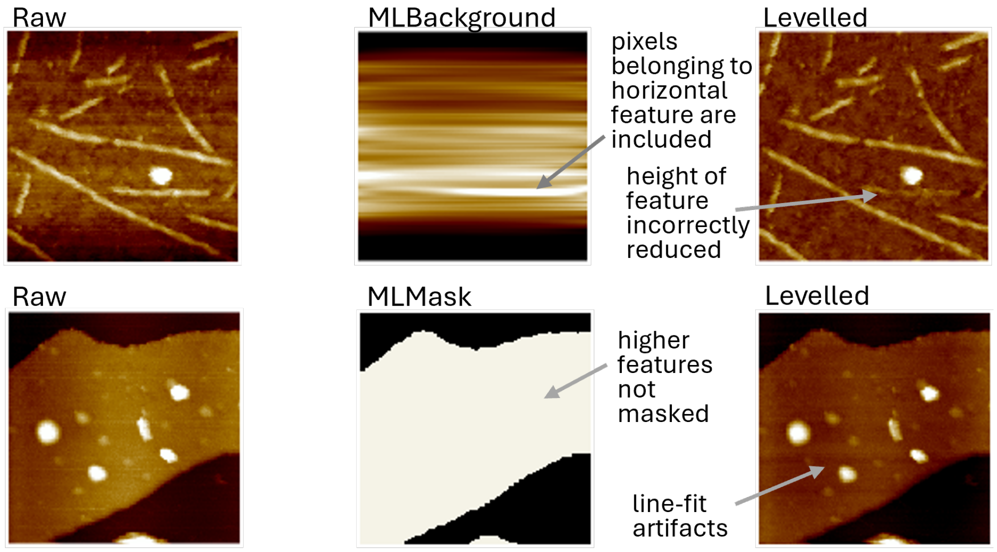


Figure S8. Examples of two flaws in the models: detection of horizontal fibrous features as noise in MLBackground, and the inability of MLMask to detect both high and low features.
